# Sexual dimorphism in adult zebrafish marrows regulates blood cell composition and immune response

**DOI:** 10.64898/2026.09.24.754168

**Authors:** Shannon E. Paquette, Carolyn L. Winston, Elizabeth A. Jones, Dionna M. Kasper

## Abstract

Hematopoiesis sustains lifelong blood production and immunity and is strongly influenced by biological sex. Males and females differ in their hematopoietic composition and susceptibility to infection and hematologic disease. Because current models cannot readily disentangle the independent contributions of sex chromosomes versus hormones, the mechanisms underlying sexual dimorphism remain poorly understood. Zebrafish have proven to be a powerful, translatable system for studying hematopoiesis, and offer a unique opportunity to isolate hormonal contributions due to their absence of sex chromosomes. However, whether sex is regulated in the zebrafish hematopoietic system remained unknown. Here, we discovered pronounced sexual dimorphism in the adult zebrafish marrow, with compositional differences peaking at reproductive age. Transcriptome analysis of marrows revealed conservation of sex-specific programs between zebrafish and human datasets. Upon immune challenge with lipopolysaccharide, all major blood lineages exhibited sex-biased changes in their composition and transcriptomes with the largest responses in erythroid and myeloid cell types. Among these sex differences was an unexpected immunoregulation of the erythroid lineage across maturation states. These findings establish biological sex as a major regulator of zebrafish hematopoiesis and immunity, paralleling humans, and lay the groundwork necessary for future mechanistic studies on sexual dimorphism in hematologic health and disease.

## INTRODUCTION

Hematopoiesis is the dynamic and tightly regulated process that supports blood cell production and immunity throughout life. Biological sex is recognized as a fundamental regulator of mammalian hematopoiesis, influencing hematopoietic stem and progenitor cell (HSPC) lineage output^1–3^, proliferative capacity^4^, transplantation success^5–7^, and leukemic potential^5,8^. Sexual dimorphism extends to downstream differentiated hematopoietic lineages as well, with males and females exhibiting distinct immune cell compositions and functional responses to inflammation or immune challenge^3,9–11^. Importantly, these sex-driven immune biases are implicated in disease susceptibility and treatment outcomes^12^, spanning infectious, autoimmune, and malignant diseases^13–15^. For example, approximately 8-10% of the population have an autoimmune condition, with women accounting for nearly 78% of affected individuals^13^. Conversely, males exhibit a higher incidence of most hematologic malignancies, including acute myeloid leukemia and multiple myeloma, and frequently experience poorer clinical outcomes^15^. As such, incorporating biological sex as a variable across hematologic studies is imperative to elucidating both homeostatic and disease-associated mechanisms regulating blood cell function.

Although biological sex is a critical determinant of hematopoietic regulation, little is known about the precise cellular and molecular mechanisms underlying these differences. Sexual dimorphism in the hematopoietic system is influenced by both sex chromosome complement (e.g., XX vs XY)^16,17^ and steroid hormone signaling^4^; however, disentangling the relative contributions of these intrinsic genetic factors versus extrinsic hormonal influences on blood cell composition and function is complex^18,19^. The X chromosome harbors a greater number of immune-related genes, many of which are hormonally regulated or escape X-inactivation^20,21^. Divergent hormone signaling also contributes to differing hematopoietic function. Estrogen biases responses toward humoral immunity, increasing female susceptibility to autoimmune diseases, whereas testosterone often suppresses adaptive responses^21^. Moreover, hormone levels vary throughout life, and transient fluctuations can have lasting effects on the hematopoietic system^21^, further complicating our ability to isolate individual drivers of blood cell functions.

Various mouse models have been developed to disentangle the effects of sex chromosome complement from gonadal sex and to determine how chromosome dosage contributes to dimorphic phenotypes. Such models include the Four Core Genotypes and XY* mouse strains, both of which require analysis of a minimum of four different sex chromosome complements^22, 23^. However, very few studies have applied them to hematopoietic cell biology. This could be due to constraints in scalability from research costs, experimental complexity, or technical caveats of these models. These limitations highlight the need for complementary vertebrate models to identify conserved mechanisms underlying sexual dimorphism in the hematopoietic system.

The zebrafish (*Danio rerio*) model provides a powerful and underutilized opportunity for addressing sex differences in hematopoietic composition and immunity. They possess all the major blood cell types, immune functions, and equivalent hematopoietic organs as mammals, and their hematopoietic regulatory pathways are highly conserved^24–26^. Zebrafish have long been lauded for their contributions to defining mechanisms of hematopoiesis and have proven to be an efficient and translatable system^27–34^. Zebrafish are used to model viral and bacterial virulence^35,36^. Expression of human disease-causing genes or mutations in their orthologs in zebrafish recapitulate human disease phenotypes, including various types of leukemia, myeloproliferative neoplasms (MPN), and immune-mediated inflammatory diseases such as inflammatory bowel disease^32,37–43^. Zebrafish studies have also led to the identification of new therapeutic targets. The drug 16,16-dimethyl-Prostaglandin E2 was first discovered in zebrafish to improve HSPC engraftment and was later approved for clinical trials of bone marrow transplantation in human patients^32–34^. Notably, laboratory zebrafish strains lack heteromorphic sex chromosomes and rely on polygenic mechanisms, environmental conditions, and hormones to determine their sex^44,45^. This positions zebrafish as a novel model to dissect endocrine control of sex-specific hematopoietic function with the absence of chromosomal influence. However, the role of biological sex on their hematopoietic systems remains undefined.

Here, we determine that biological sex influences hematopoietic composition and function in the adult zebrafish kidney marrow, the functional equivalent of the mammalian bone marrow^46^. We characterize the cellular and transcriptional profiles of hematopoietic cells from all major lineages and the surrounding microenvironment niche in male and female marrows under homeostatic and inflammatory conditions. Through comparative analyses with human marrow datasets, we also reveal several sex-specific transcriptomic signatures shared between the species, supporting zebrafish as a translatable model to uncover mechanisms of sexual dimorphism in the hematopoietic system. Lastly, we demonstrate that sex differences are pervasive across hematopoietic lineages. Erythroid and myeloid lineages showed the greatest number of sex-biased changes in response to an inflammatory stimulus, with unique expression dynamics across their maturation states between sexes. Together, this work further strengthens zebrafish as a complementary vertebrate model for investigating mechanisms regulating adult hematopoiesis and underscores the importance of incorporating biological sex into hematologic studies.

## RESULTS

### Adult zebrafish marrow composition varies by sex and age

We investigated whether biological sex shapes hematopoietic composition in adult zebrafish by conducting a longitudinal analysis of marrows from males and females from 2 months of age (juvenile stage) through 24 months of age (late adult stage) using flow cytometry (**Fig. 1**). Using a well-established gating strategy^27,37,47^, we quantified the percentage of live, single cells within four broad gates, which are composed mainly of precursor, myeloid, erythroid, or lymphoid cell types (**Fig. 1a, b**). Strikingly, hematopoietic composition was temporally regulated and distinct between males and females at specific ages during their lifespan (**Supplementary Fig. 1a, b, Supplementary Table 1**). Sex-specific differences in the myeloid, erythroid, and lymphoid gates began at the onset of sexual maturity at 3 months of age and became most pronounced at 6 months of age, during peak reproductive capacity^48^ (**Fig. 1d-f**). These findings reveal that both sex and age are key determinants of zebrafish marrow composition and uncover a previously unappreciated period of heightened hematopoietic divergence at 6 months old.

**Fig. 1:**
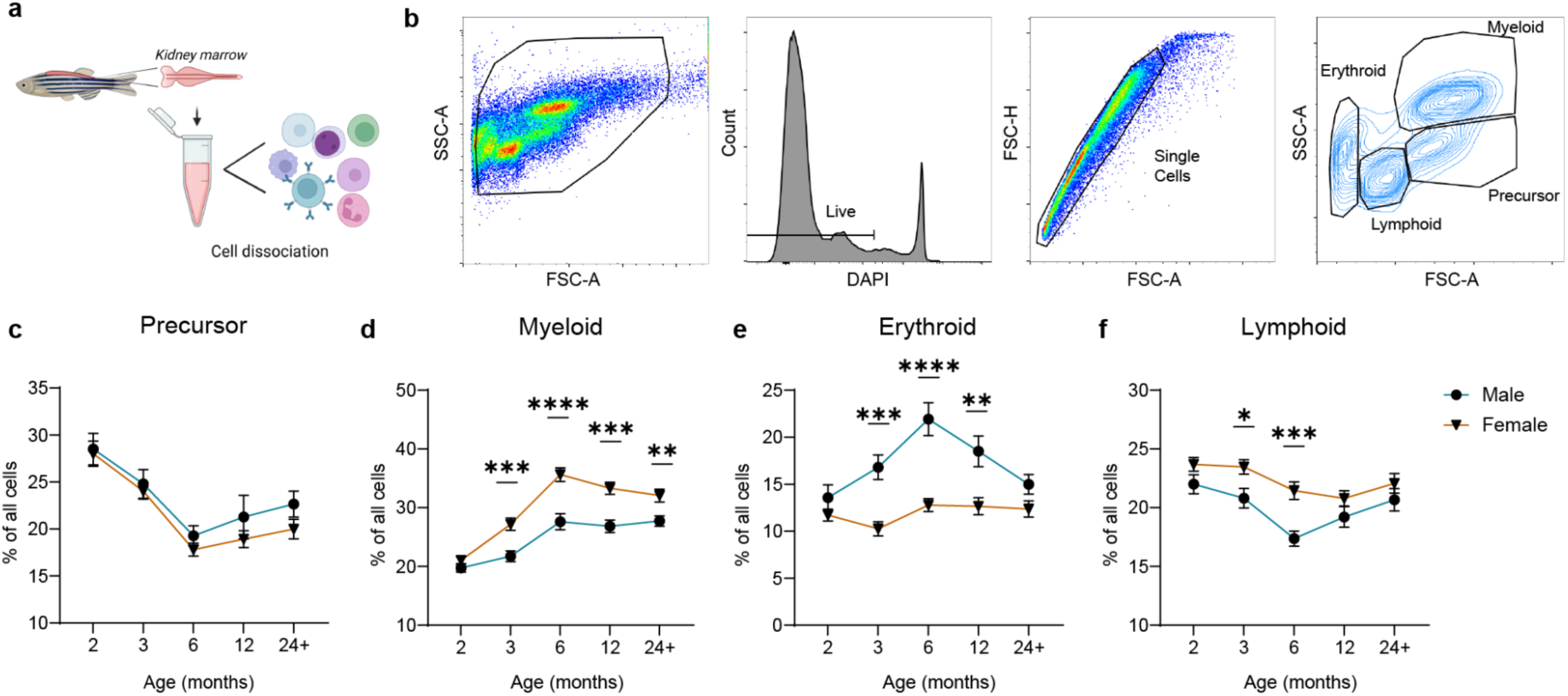
Male and female zebrafish have distinct hematopoietic composition within the marrow. (a) Diagram of kidney marrow dissection and single cell dissociation for flow cytometric analysis. (b) Representative gating strategy used to examine hematopoietic composition in adult zebrafish marrow. (c-f) Plots represent the percent of total cells in the marrow that fall within each hematopoietic gate: (c) precursor, (d) myeloid, (e) erythroid, and (f) lymphoid. Ages range from 2 to 24+ months old. n = 15-20 animals per group. pval <0.05 Significance was determined using an unpaired t-test with Welch’s correction between males (blue lines, circles) and females (orange lines, triangles) at each designated time point. Error bars represent SEM.

### Sex-specific transcriptional programs are conserved across vertebrates

We tested whether marrows were also sexually dimorphic at the transcriptional level by performing bulk RNA-sequencing (RNA-seq) on male and female marrows at 6 months of age when sex differences in hematopoietic composition were the greatest (**Fig. 1d-f**). We conducted differential gene expression analyses utilizing DESeq2^49^ and uncovered 1,073 significant differentially expressed genes (DEGs) across the sexes (**Fig. 2a, Supplementary Table 2**). Gene ontology (GO) term enrichment analysis of male- and female-specific DEGs uncovered both shared and distinct categories among the top 20 most significant GO terms (**Fig. 2b, Supplementary Table 3**). Marrows from both sexes were enriched in pathways relevant to blood cell biology and hormone signaling. However, females showed greater enrichment for immune functions, with 4 GO terms representing 18 unique DEGs, whereas males had only 1 hematopoietic GO term comprising 7 DEGs (**Fig. 2b, Supplementary Table 3**). GO categories unique to males were associated with metabolism and intracellular transport, while terms associated with cell cycle and reproductive processes predominated in females. Thus, these findings demonstrate that male and female zebrafish marrows are transcriptionally distinct at steady-state.

**Fig. 2:**
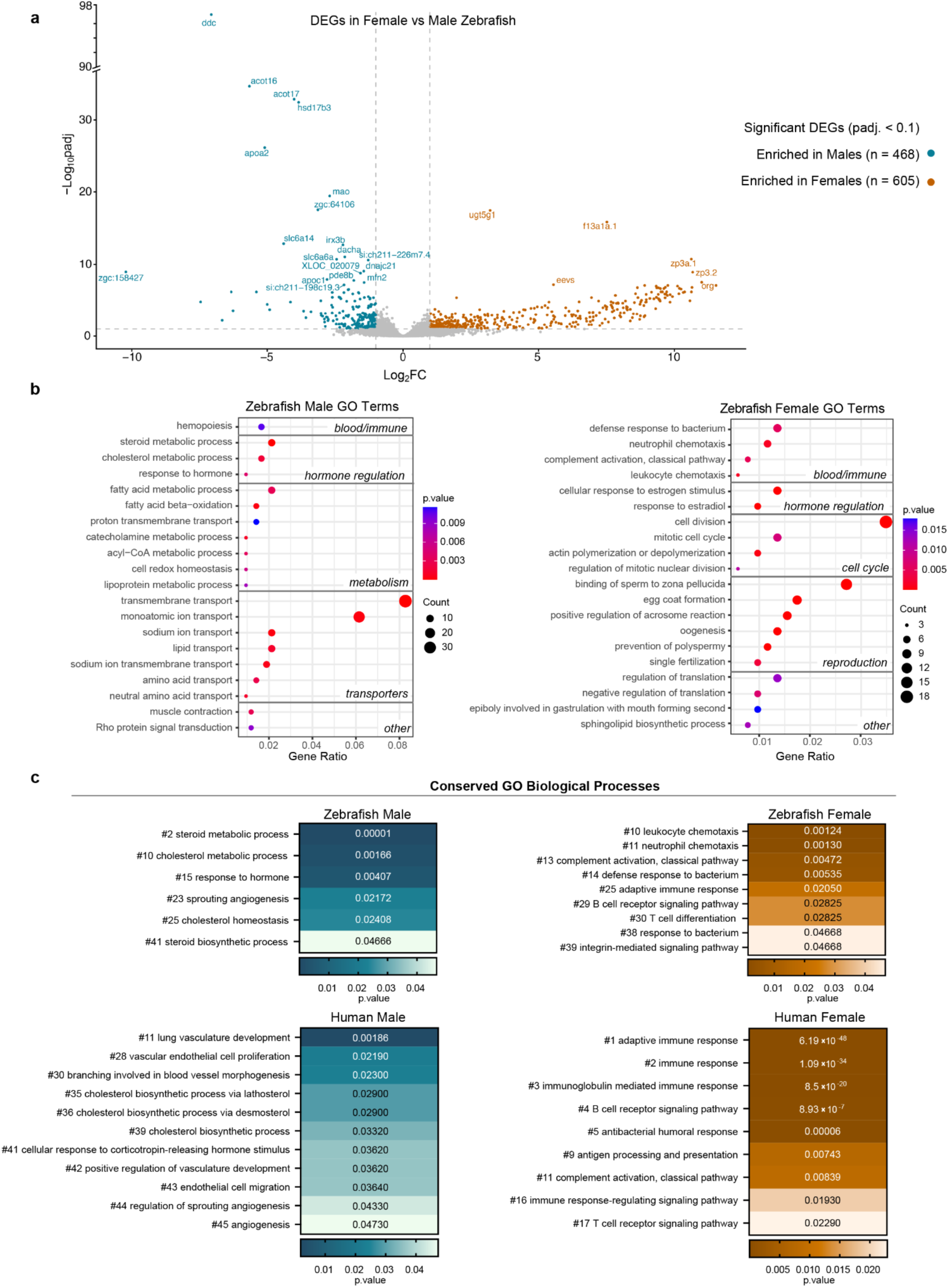
Male and female zebrafish have distinct transcriptional profiles at 6 months old. (a) Volcano plot of bulk RNA-seq data of male and female marrows from PBS-injected zebrafish (n = 5 marrows per sex). Genes with significantly different expression between sexes are colored in blue or orange for males or females, respectively. padj. <0.1 and Log_2_FC <-1 or >1. Significance was determined using a Wald test. (b) Top 20 most significant biological processes enriched within zebrafish male and female marrows as determined by GO analysis. (c) GO analysis of DEGs from zebrafish and human marrow bulk RNA-seq datasets. Only related terms shared between the species are displayed. GO term number (#) represents its rank among significant terms (pval <0.05, Fisher’s Exact test).

To determine whether the biological processes that differed between zebrafish male and female marrows were conserved in humans, we analyzed a publicly available bulk RNA-seq dataset of healthy human bone marrow^50^. We performed GO enrichment analysis on human male- and female-specific DEGs and compared the significantly enriched terms to those in our zebrafish marrow dataset (**Fig. 2c, Supplementary Table 3**). Notably, many sex-specific terms were common to both zebrafish and human marrows. Male zebrafish and human samples shared processes related to angiogenesis and metabolism of steroids and the steroid hormone precursor cholesterol, whereas females in both species shared pathways related to immune function, such as bacterial response, B- and T-cell related adaptive immunity, and complement activation (**Fig. 2c**).

To identify conserved regulatory mechanisms governing sex differences within zebrafish and human marrows, we compared sex-specific DEGs for both species using Qiagen’s Ingenuity Pathway Analysis (IPA) platform^51^ (**Supplementary Fig. 2a**). Specifically, we conducted an analysis that compared enriched biological functions identified from the DEGs, as well as an analysis that predicted shared upstream regulators of the DEGs. IPA for “Biological Functions” that were in common between zebrafish and human marrows revealed processes involved in cell proliferation and survival, immune migration and trafficking, and immune cell development and homeostasis, consistent with our GO enrichment analysis (**Supplementary Fig. 2b and Fig. 2c**). IPA for shared endogenous or exogenous “Upstream Regulators” identified factors with strong, direct immunomodulatory functions, including the bacterial pro-inflammatory signal lipopolysaccharide (LPS), corticosteroid anti-inflammatory drugs methylprednisolone and dexamethasone, and various cytokines, such as tumor necrosis factor (TNF), interferon gamma (IFNG), and interleukins (IL) 2 and 4 (**Fig. 3a**). Together, these analyses identify sex-specific transcriptional programs and upstream regulatory factors that are conserved between zebrafish and human marrows, particularly in pathways related to homeostatic immune function and inflammatory signaling.

**Fig. 3:**
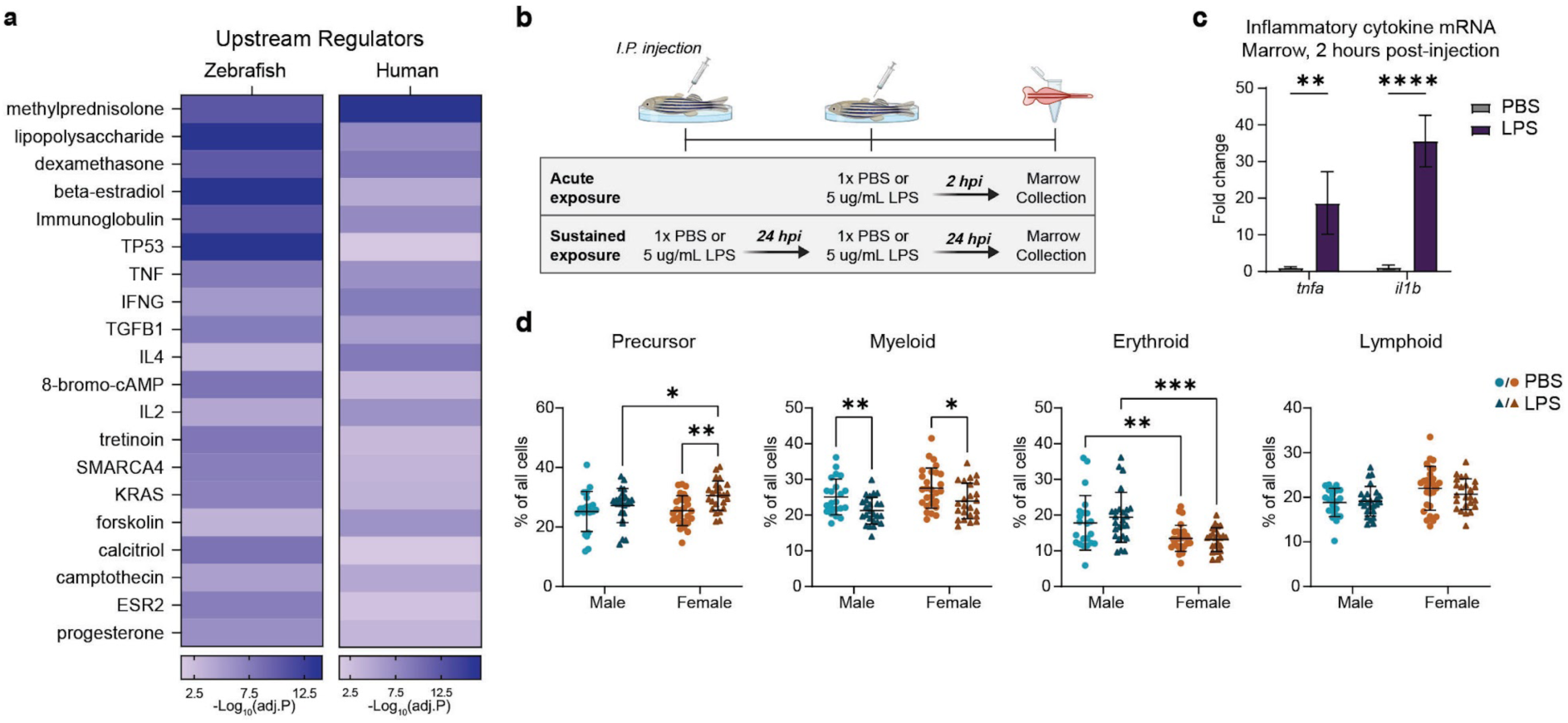
LPS exposure induces sex-biased change in hematopoietic composition in the marrow. (a) IPA for predicted upstream regulators shared between zebrafish and human marrows derived from bulk RNA-sequencing datasets (padj. <0.1, Fisher’s Exact test). (b) Diagram of PBS (control) and LPS exposure paradigm by intraperitoneal injections of 6 month old zebrafish. Marrows were harvested 2 hours post injection (hpi) to assess acute responses via qRT-PCR or bulk RNA-seq analysis, or after 2 consecutive injections, 24 hpi apart for compositional analyses of sustained exposure. (c) *tnfa* and *il1b* transcript levels in the marrow 2 hpi of PBS- or LPS-treated zebrafish. n=3-4 per group. Unpaired t-test. (d) Plots represent the percent of total cells in the marrow that fall within each broad hematopoietic gate: precursor, myeloid, erythroid, and lymphoid for PBS- and LPS-treated males and females. n= 22-27 per group. pval <0.05, Two-Way ANOVA.

### Inflammatory challenge elicits a sex-biased immune response

Both GO analysis and IPA strongly suggested that inflammation would induce a sex-specific immune response. Therefore, we challenged 6 month old male and female zebrafish with LPS, a top hit in the IPA for Upstream Regulators (**Fig. 3a**). To elicit a biologically relevant inflammatory response, we injected sub-lethal doses of LPS or sterile PBS into the peritoneal cavity (**Fig. 3b**). Expression analysis of the pro-inflammatory cytokines *tnfa* and *il1b* confirmed a strong inflammatory response to acute LPS exposure in the marrow within 2 hours post injection (hpi) (**Fig. 3c**). To allow time for cellular composition to shift in response to LPS exposure, we treated zebrafish for two consecutive days and performed flow cytometry analysis on the following day (i.e., sustained exposure) (**Fig. 3b**). The zebrafish marrow responded in similar ways to mammals after sustained LPS exposure. In mice, LPS induces trafficking of mature myeloid cells out of the marrow, and those cells are then subsequently replenished by HSPC proliferation^52^. Indeed, zebrafish also showed a reduction of myeloid cells following LPS exposure (**Fig. 3d**). Interestingly, we observed a sex-biased immune response in the precursor population, in which the proportion of cells in the precursor gate was significantly increased upon LPS exposure only in female marrows (**Fig. 3d**). In contrast, neither males nor females had changes in the lymphoid or erythroid gates following LPS treatment (**Fig. 3d**). These results show hematopoietic composition responds to immune challenge in sex- and cell type-dependent ways.

Compositional differences following sustained LPS exposure may be driven by divergent initiators of response pathways between males and females. Therefore, to understand the early transcriptional responses following acute inflammatory challenge, we conducted bulk RNA-seq on male and female marrows collected two hours following intraperitoneal injection of LPS or PBS (i.e., acute exposure) (**Fig. 3b**). Independent analysis of male and female datasets revealed a total of 151 genes with significantly altered expression following acute LPS exposure (**Fig. 4a**). This list of LPS-responsive genes is 2.3 times greater in number than if the datasets from both sexes were combined (i.e., 151 versus 66 genes) (**Fig. 4a**). Of the 151 genes, 95% had altered expression upon LPS challenge in only one of the sexes, whereas just 7 LPS-responsive genes were in common between the sexes (*mxf*, *si:dkey-195m11.11*, *tlr5b*, *si:ch211-147m6.1*, *gpr84*, *ppm1j*, *rhogb*) (**Supplementary Fig. 3**). Notably, the identification of sex-specific expression dynamics of many LPS-responsive genes would have been missed if sex was disregarded as a biological factor (**Supplementary Table 4**).

**Fig. 4:**
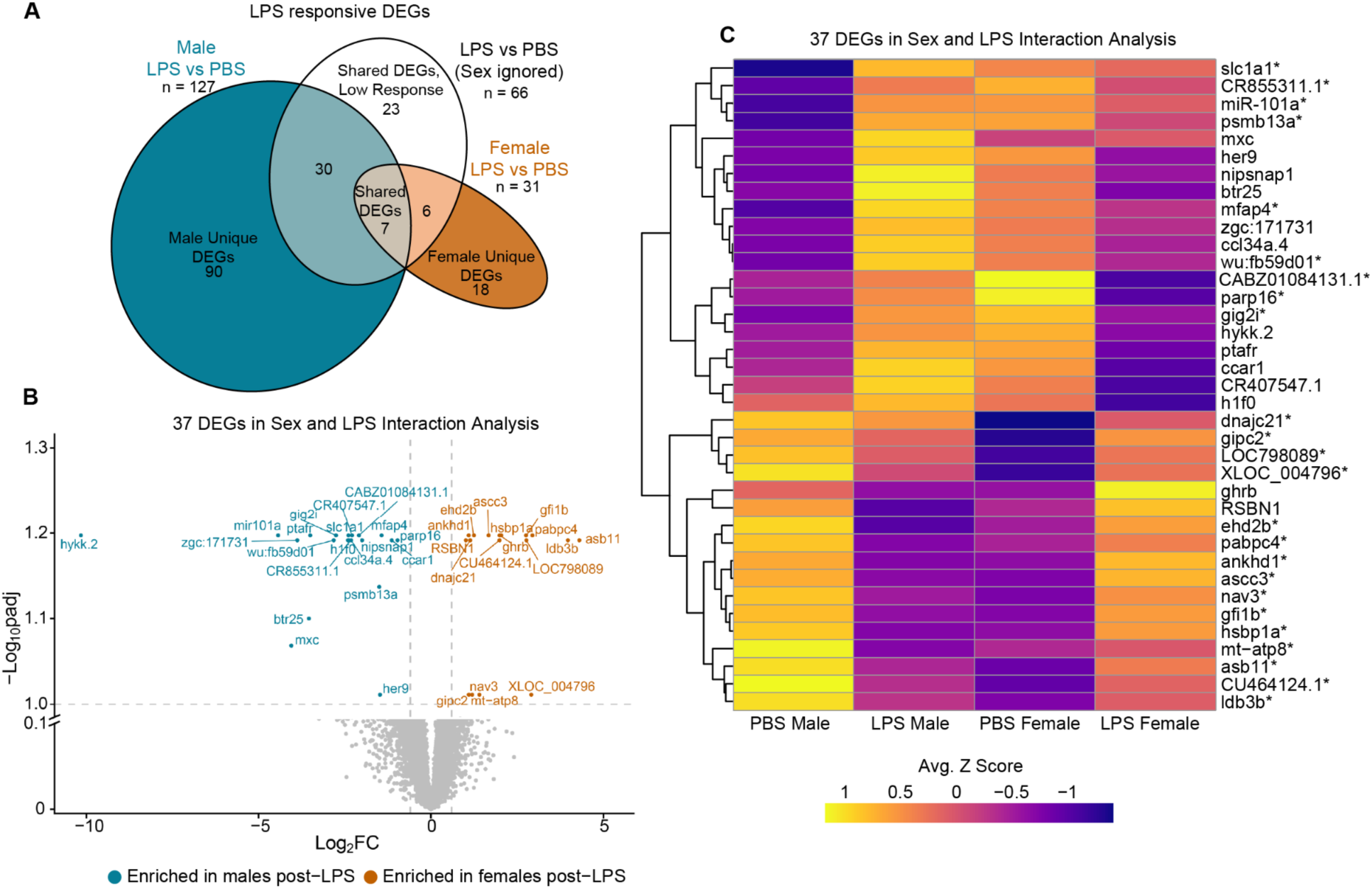
Transcriptional responses to LPS exposure in the marrow are dependent on sex. (a) Venn diagram shows overlap of DEGs following bulk RNA-seq of 6-month old marrows from fish exposed to LPS or PBS. Groups in the diagram correspond to DEG analyses of each sex individually or the sexes combined. n= 5 per group. padj. <0.1. (b) Volcano plot of 37 significant DEGs from interaction analysis of sex and LPS in marrows. padj. <0.1 and Log_2_FC >0.6 or <-0.6. Wald test used to calculate significance. (c) Heatmap of averaged Z score of 37 DEGs from sex and LPS exposure interaction. *Genes represent DEGs that were also differentially expressed between PBS Female and PBS Male.

We next identified which genes respond to LPS in a sex-dependent manner by conducting an interaction analysis, which accounts for baseline sex differences in expression in our bulk RNA-seq dataset. We found 37 genes responded to acute LPS exposure in a sex-dependent manner (**Fig. 4b, c**). Interestingly, many genes that were differentially expressed between sexes in the control group exhibited inverse expression patterns following the LPS challenge (**Fig. 4c**). These data highlight that early transcriptional responses to an LPS-induced inflammatory challenge are highly dependent on sex.

### Single cell RNA sequencing identifies male and female cell type-specific changes in composition following inflammatory challenge

A variety of cell types reside in the whole kidney marrow, including hematopoietic cells from all the lineages, vascular cells, and epithelial cells, which support filtration functions of the kidney^53^. Thus, the sex biases observed in bulk tissue analyses could be due to broad lineage shifts, changes in proportions of specific cell types or subpopulations of those cell types, and/or transcriptional differences within the same or different cell types. So, to understand the cellular basis of sex differences in the hematopoietic system, we conducted single cell RNA-sequencing (scRNA-seq) of marrows from 6 month old male and female zebrafish injected with sublethal doses of LPS or the PBS control. Using the same sustained LPS exposure paradigm as for flow cytometry analyses, treatments occurred on 2 consecutive days to allow for the marrow to respond both transcriptionally and compositionally to the inflammatory challenge (**Fig. 3b**). Dissected marrows underwent scRNA-seq using the 10X Genomics Chromium system, followed by the removal of low-quality cells and doublets from the resulting datasets. We conducted unsupervised clustering of merged data sets using Seurat and used known marker genes to identify 32 distinct clusters. These clusters captured the entire suite of hematopoietic lineages along with other cells of the kidney^53^ (**Fig. 5a, Supplementary Table 5**).

**Fig. 5:**
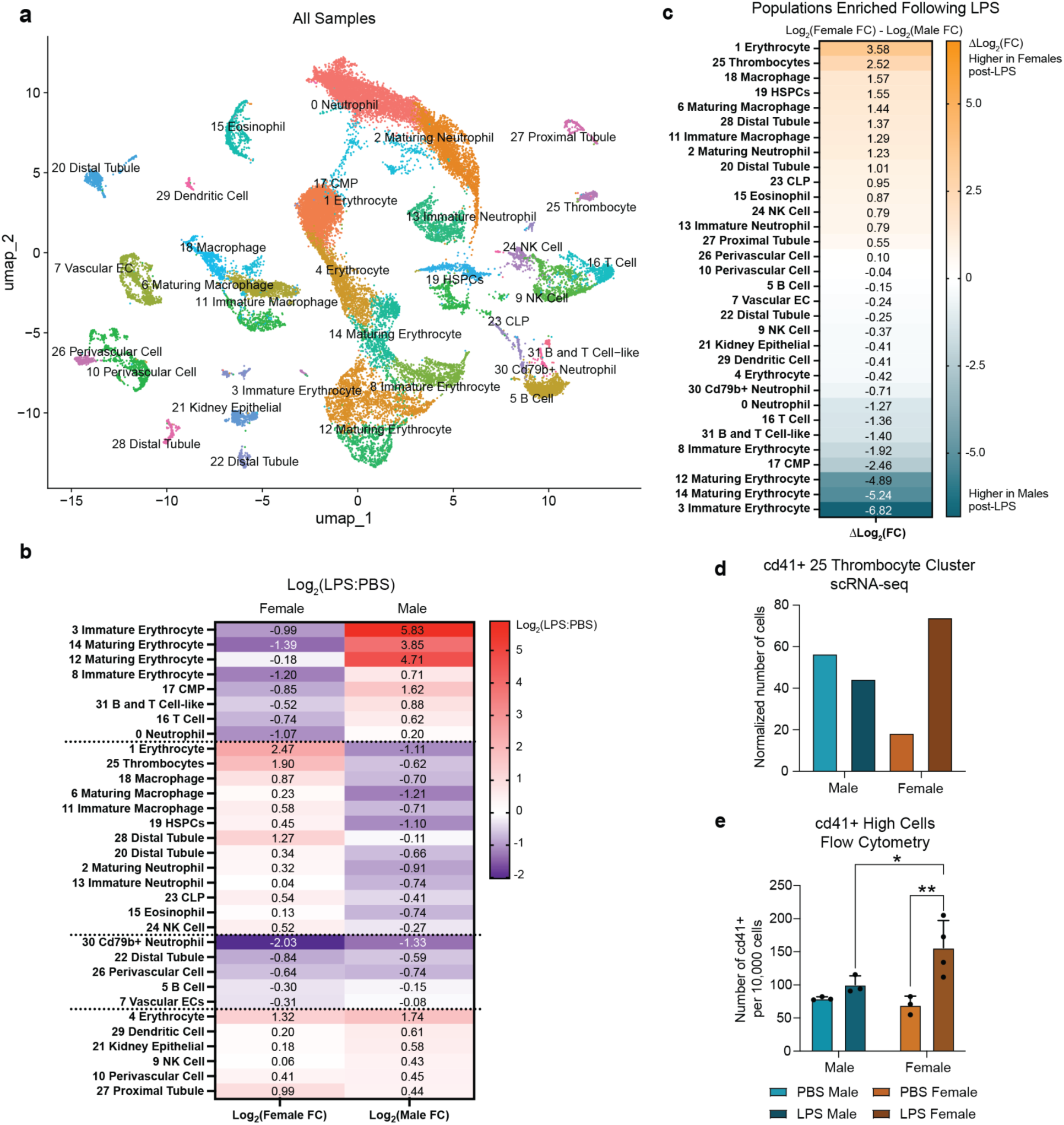
scRNA-seq reveals sex-specific compositional responses to LPS across hematopoietic lineages. (a) Annotated Uniform Manifold Approximation Projection (UMAP) representing all 32 captured cell populations within 6 month old marrows across 4 groups: PBS male, LPS male, PBS female, LPS female. Fish were injected with PBS or LPS for 2 consecutive days and collected for single cell RNA-sequencing 24 hpi on the third day. (b) Log_2_ fold change of normalized cell numbers in LPS vs PBS samples per cluster for males and females. (c) Differential of the Log_2_FC displayed in (b). (d) Number of *cd41* transcript+ cells (i.e., *itga2b+*) within the 25 Thrombocyte cluster of the scRNA-seq dataset, normalized to the total number of all cells per sample. (e) Number of *Tg(cd41:GFP)*+ high cells within the marrow per 10,000 cells determined by flow cytometry analysis. n= 3-4 per group. Two-Way ANOVA.

With the enhanced cellular resolution of the scRNA-seq dataset, we found that sexually dimorphic responses to inflammatory challenge were not just limited to precursor populations as observed with flow cytometry analysis, but were pervasive throughout the marrow (**Fig. 5b, Supplementary Fig. 4a, b**). More than half of the cell clusters (17/32) had a greater than 2-fold differential in abundance between the sexes following sustained LPS treatment (**Fig. 5c**), including progenitor populations such as cluster 19, HSPCs; cluster 23, Common Lymphoid Progenitor (CLP); and cluster 17, Common Myeloid Progenitor (CMP). The most dramatic compositional changes occurred within the erythroid lineage clusters (**Fig. 5b, c**). The abundance of *itga2b/cd41*+ thrombocytes (cluster 25) (**Fig. 5d**) and other mature erythroid clusters (clusters 1 and 4) were increased in female marrows upon LPS treatment, whereas the less mature erythroid populations (clusters 3, 8, 12, 14) were expanded in exposed male marrows (**Supplementary Fig. 4c**). Consistent with these observations, the number of thrombocytes, marked by high fluorescence of *Tg(itga2b/cd41:GFP)*^54^, were significantly increased in females following LPS exposure via flow cytometry analysis (**Fig. 5e**).

Other clusters with a greater than 2-fold differential in cell abundance between sexes included all macrophage clusters (clusters 11, 6, 18), the more mature neutrophil clusters (clusters 2, 0), T-cell related clusters (clusters 16 and 31) and distal tubule clusters (clusters 20, 28) (**Fig. 5c**). Like the erythroid lineage clusters, LPS-induced cell abundance changes within a given lineage or cell type did not always occur in the same direction between sexes. In the myeloid lineage for example, mature neutrophils were enriched in LPS-treated males, whereas less mature neutrophils and all macrophage subtypes were enriched in LPS-treated females (**Fig. 5b, c**). When cells were combined across the more mature myeloid clusters, both sexes showed a decrease in cell abundance upon LPS exposure, like the observations made for the myeloid gate in flow cytometry analysis (**Supplementary Fig. 4d, Fig. 3d**). Thus, the sex-biased compositional changes in myeloid cell populations became masked when assessed with bulk analyses. Taken together, scRNA-seq uncovered widespread differences in marrow composition between sexes in response to inflammatory challenge, spanning all hematopoietic lineages and the microenvironment. Sexual dimorphism was greatest in erythroid populations and showed distinct patterns across maturation states and lineage-related cell types.

### scRNA-seq identifies cell type-specific changes in transcription between the sexes following inflammatory challenge

To investigate the cellular sources of the sex-specific transcriptional differences in response to LPS, we identified the genes that were differentially expressed between LPS and PBS treatments for male and female marrows across all scRNA-seq clusters. Similar to the observations made for hematopoietic composition, transcriptional differences between sexes upon LPS exposure were extensive, impacting clusters for all hematopoietic lineages, along with the structural cells of the kidney. 59% (19/32) of clusters had more than 100 DEGs following LPS exposure, with erythroid and myeloid clusters containing the greatest numbers of DEGs (**Fig. 6a**). Furthermore, the majority of DEGs in all hematopoietic clusters were unique to a sex, with only a minority of genes differentially expressed in the same directionality in both sexes (**Fig. 6a**). Interestingly, 21 out of 32 clusters had a greater number of DEGs in female marrows compared to those in males, suggestive of a more reactive immune system in female zebrafish, like female mammals^55,56^.

**Fig. 6:**
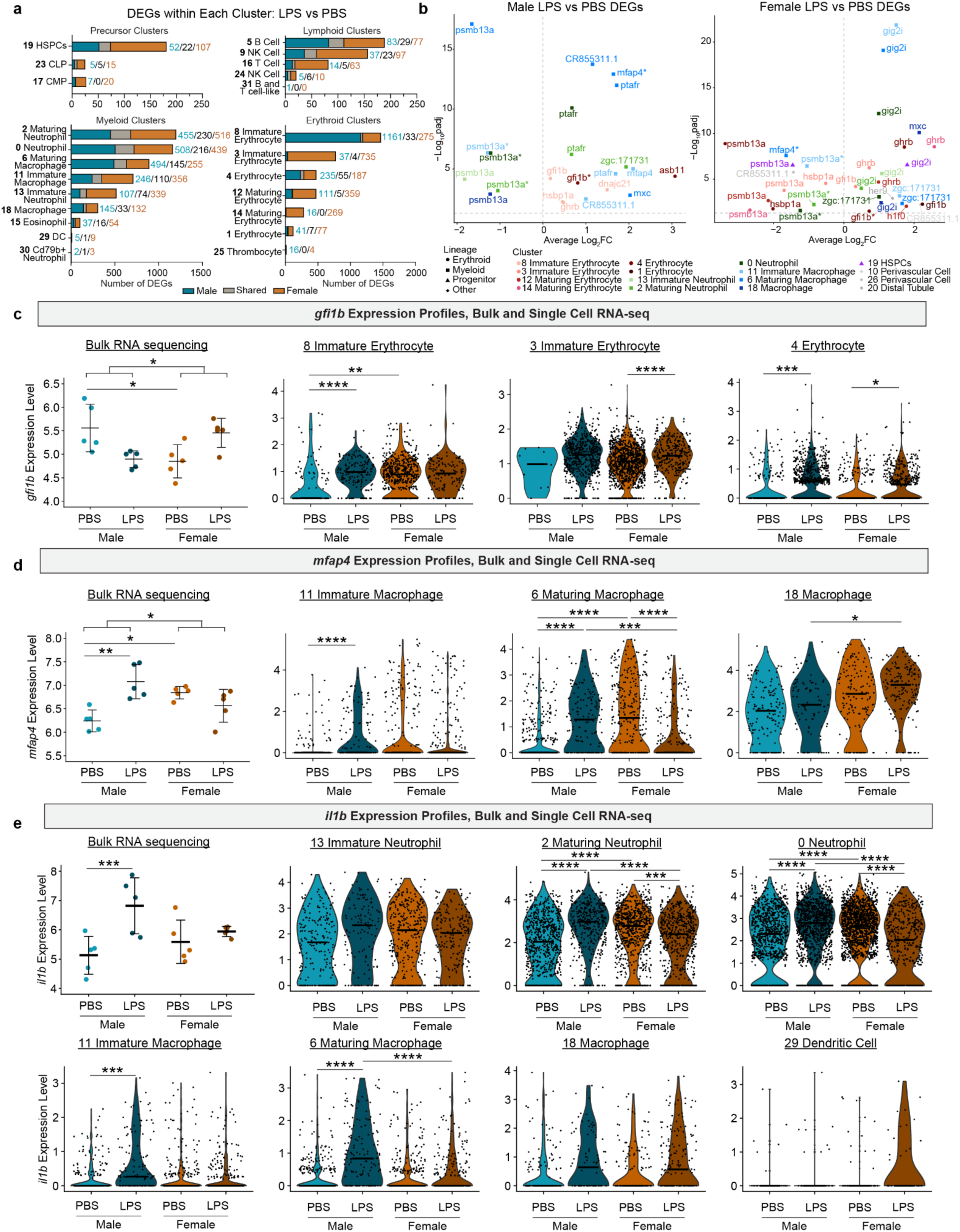
Maturation state influences sex-specific transcriptional programs at baseline and following immune challenge. (a) Number of LPS vs PBS DEGs with the same Log_2_FC directionality shared between males and females, male-unique DEGs, and female-unique DEGs in hematopoietic clusters. DEGs filtered by padj <0.05, Log_2_FC >0.25 or <-0.25, and male or female LPS/PBS cells with >10% transcript expression. (b) Volcano plot of only the LPS vs PBS DEGs in males and females with padj <0.05 and LPS/PBS cells with >10% transcript expression that also appeared as DEGs in the bulk RNA-seq interaction analysis. (c) Normalized *gfi1b* transcript expression levels in bulk RNA sequencing data, and clusters 8, 3, and 4 from single cell RNA sequencing data (padj <0.05). (d) Normalized *mfap4* transcript expression levels in bulk RNA sequencing data (padj <0.1), and clusters 11, 6, and 18 from single cell RNA sequencing data (padj <0.05). (e) Normalized *il1b* transcript expression levels in bulk RNA sequencing data (padj <0.1), and clusters 13, 2, 0, 11, 6, 18, and 29 from single cell RNA sequencing data (padj <0.05). Wald test used to calculate significance for bulk RNA-sequencing data. Wilcoxon test used to calculate significance for single cell RNA-sequencing data.

Next, we analyzed the distribution of expression across scRNA-seq clusters for genes that responded in a sex-dependent manner to acute LPS exposure in our bulk RNA-seq dataset (**Fig. 4b, c**). We detected expression for 35 out of the 37 of these genes. All hematopoietic clusters expressed at least one of the 35 detectable genes in at least one condition, and many genes were present in more than one hematopoietic cluster (**Supplementary Fig. 4e**). Analysis of the 35 genes for significant differences in gene expression following sustained LPS exposure in each sex identified 55 DEGs (as some genes were expressed in more than one cluster), many of which had cluster- and sex-specific changes (**Supplementary Table 6**). In male marrows, 22 DEGs, consisting of 11 unique genes, changed across different clusters from the erythroid and myeloid lineages (**Fig. 6b, Supplementary Table 6**). In female marrows, 33 DEGs, encompassing 11 genes that partially overlapped the male set, changed in erythroid and myeloid clusters as well as within the HSPC and three non-hematopoietic clusters (**Fig. 6b, Supplementary Table 6**). Interestingly, 82% (45/55) of DEGs had expression patterns that were unique to a sex in a given cluster. For example, *platelet-activating factor receptor* (*ptfar*) was only significantly changed in myeloid clusters and in LPS-treated males. Similarly, *growth hormone receptor b* (*ghrb*) was significantly altered in both LPS-treated males and females, but in different erythroid clusters (**Fig. 6b**). Only three genes showed differential expression upon LPS exposure in both sexes across five clusters, namely *proteasome 20S subunit beta 13a* (*psmb13a*)*, growth factor independent 1B transcription repressor* (*gfi1b*), and *microfibril associated protein 4* (*mfap4*) (**Fig. 6b**). These data indicate that sex-specific transcriptional responses to inflammatory challenge occur within particular hematopoietic cell types and are likely a contributing factor to the sexual dimorphic response which was first observed in the bulk RNA-seq data.

### Hematopoietic subpopulations have unique sex-specific transcriptional responses to inflammatory challenge

The observation that LPS exposure altered gene expression in specific clusters prompted us to analyze transcriptional patterns across subpopulations of an individual hematopoietic cell type in our scRNA-seq dataset. Given that the greatest immune responses to LPS occurred in the erythroid and myeloid populations, we focused on genes that are well known regulators of these lineages. *Gfi1b* functions in erythrocyte differentiation^57^ and was one of the genes that showed a sex-specific transcriptional response to acute LPS exposure in our bulk RNA-seq dataset (**Fig. 4b, c**). In the scRNA-seq dataset, expression of *gfi1b* was enriched in clusters of the erythroid lineage (**Supplementary Fig. 5a, b**) but surprisingly had sex-specific dynamics which varied across erythroid subpopulations upon sustained LPS exposure. The early upregulation of *gfi1b* in whole female marrows persisted with the sustained LPS treatment and occurred only in mature clusters 1 and 4 and immature erythroid cluster 3 (**Fig. 6c**). In male marrows, the early decline of *gfi1b* expression from acute LPS exposure then increased significantly in a different set of erythroid clusters, clusters 4 (mature) and 8 (immature), during sustained exposure (**Fig. 6c**). Notably, some erythroid clusters did not have differences in transcript levels between groups, including the maturing erythroid clusters 12 and 14, and thrombocyte cluster 25 (**Supplementary Fig. 5c**). Thus, the complex dynamics of *gfi1b* expression in the erythroid clusters demonstrate that sex-biased responses depend on particular erythroid subpopulations of differing maturation states.

Similarly, *mfap4*, a regulator of myeloid differentiation^58^ and a sexually dimorphic LPS-responsive gene in the bulk RNA-seq dataset, had varied expression across macrophage subpopulations in the scRNA-seq dataset (**Supplementary Fig. 5a, b**). Like the early transcriptional response to acute LPS treatment (**Fig. 4b, c**), sustained LPS exposure increased *mfap4* expression in males, but only in the less mature macrophages, clusters 11 and 6 (**Fig. 6d**). In female marrows, *mfap4* expression changed in the opposite direction to males and was significantly downregulated only in the maturing macrophages cluster 6, consistent with the downward trend seen in the acute LPS exposure (**Fig. 6d**). While the less mature macrophage clusters (clusters 11 and 6) clearly showed sex-biased responses to LPS, mature macrophages (cluster 18) showed no large differences in *mfap4* transcript levels between groups, suggesting that maturation state could play a role in sex-biased responsiveness to inflammatory challenge.

We also examined the expression dynamics of *il1b*, an important pro-inflammatory cytokine known to be responsive to LPS^59^. *Il1b* expression was enriched across clusters in the myeloid lineage including the common myeloid progenitor, dendritic cells, and all macrophage and neutrophil subtypes (**Supplementary Fig. 5a, b**). Again, we find that sex-specific expression patterns were distinct across subpopulations of a given cell type. For neutrophil subtypes, *il1b* expression was upregulated in LPS-treated males compared to a downregulation in LPS-treated females, which occurred only in more mature clusters (clusters 2 and 0) (**Fig. 6e**). No significant differences in transcript levels appeared in immature neutrophils (cluster 13) for any group. Interestingly, for some of the macrophage subpopulations, sex-specific changes in *il1b* expression were evident only in the less mature clusters (clusters 11 and 6). While macrophages had a similar decrease in *il1b* expression in LPS-treated males as neutrophils, transcript levels changed in alternate directions in LPS-treated females between these myeloid lineage cell populations (**Fig. 6e**). Thus, the strong upregulation of *il1b* in LPS-treated males compared to a weaker one in females in the bulk RNA-seq is illuminated by the congruent and contrasting expression patterns in neutrophil and macrophage clusters for the respective sexes. Taken together, analysis of *gfi1b, mfap4,* and *il1b* expression patterns in the scRNA-seq dataset reveal that sex-specific responses to LPS occur not only in particular cell types, but by specific subpopulations of these cell types, such as maturation state.

## DISCUSSION

Historically, biomedical research has been heavily male-biased across the entire translational spectrum—from cell culture to animal models to clinical trials^60^. These biases extend to the field of hematopoietic cell biology and have contributed to disparities in our understanding of hematologic disease mechanisms and therapeutic responses in males versus females^61,62^. With a pressing need to include both sexes in experimental designs, policy changes have been implemented, but only recently^63^. Consequently, most published studies on hematopoiesis and immunology in adult zebrafish either did not report biological sex or used mixed-sex cohorts without stratifying analyses by sex^53,59,64,65^. Given the increasing use of adult zebrafish to model and treat human diseases^32,66^, including many with an inflammatory component^67^, this study sought to systematically investigate sex-based differences in adult zebrafish hematopoiesis and immune response. Here, we discover widespread sexual dimorphism in the marrow, spanning all hematopoietic lineages and the surrounding microenvironment. These previously unrecognized sex differences occur at the cellular and molecular levels, and under steady-state and inflammatory conditions.

Longitudinal analysis of the four major hematopoietic lineages by flow cytometry showed that under homeostatic conditions, hematopoietic composition is dynamic in males and females throughout adulthood. Differences in the cellular makeup between sexes was most prominent in the more differentiated fractions (i.e. myeloid, lymphoid, and erythroid gates) and started at the onset of sexual maturity at three months of age. Sexual dimorphism was greatest at six months of age during the height of reproductive capacity, mirroring the dynamics of gene expression differences between males and females in five organ systems across six different species from chickens to humans^68^. Our data add another species and tissue to a growing record of evidence for a conserved developmental program that establishes sexually-dimorphic traits once an organism reaches reproductive age.

We found previously unappreciated differences in immune response between male and female zebrafish upon sub-lethal exposure to the bacterial endotoxin LPS. Bulk transcriptome profiling of marrows after an acute LPS treatment identified more than a hundred responsive genes when males and females were stratified—twice as many genes as when sexes were combined. Nearly all these LPS-responsive genes showed significant differential expression in only male or female marrows and only a handful of genes changed equivalently between the sexes. Thus, the immediate transcriptional effects of LPS-induced inflammation are highly dependent on biological sex. While we only tested LPS as a form of inflammatory challenge, these LPS-induced expression changes in zebrafish correlate with a variety of insults in humans and mice, such as sepsis, burns, and blunt trauma^69^, suggesting other immune-related transcriptional networks are likely to be tailored to a sex across species.

Sexual dimorphism in the marrow was also evident with sustained LPS treatment. Flow cytometry showed a significant expansion of the precursor gate in female marrows; however, this fraction’s exact composition is incompletely defined, making it difficult to confirm which cell types drive its sex-biased expansion. A previous study indicates that at least some HSPCs are located within the precursor gate. In zebrafish, cells that express low levels of *Tg(itga2b/cd41:GFP)* partially fall within the precursor fraction and can reconstitute the depleted hematopoietic systems of irradiated adults, a hallmark of the self-renewal properties of HSPCs^70^. Consistent with this, our scRNA-seq dataset revealed an enrichment of female HSPC and CLP clusters post-LPS treatment. In mice, HSPCs are known to divide more frequently in females than in males during homeostasis^4^. HSPC proliferation further increased upon immune challenge with LPS to replenish immune cells that mobilized from the marrow to respond to the inflammatory insult. This study, though, did not specify the sex of the mice used^71^. Besides HSPCs, less mature myeloid cells are thought to reside in the precursor gate and thus could contribute to the increase in LPS-treated females^72^. Indeed, we find a female-biased expansion of immature and maturing cells within macrophage and neutrophil populations following sustained LPS treatment in the scRNA-seq dataset. Additional work is warranted to determine the full breadth of cell types and the proportions of those cell types that fall within the precursor gate.

For more differentiated populations, the erythroid lineage showed the greatest sexual dimorphism in our scRNA-seq dataset. In unstimulated controls, we found that the more immature erythrocyte clusters predominated the female marrow, whereas mature erythrocytes were more abundant in the male marrow. These results are congruent with mammalian physiology. Female humans and mice have greater numbers of immunoregulatory erythroid precursors, CD71+ erythroid cells, whereas males have higher levels of circulating mature erythrocytes due to the stimulatory effect of testosterone on erythropoiesis^73–76^. Upon LPS stimulation, we observed a dramatic sex-biased decline in mature erythrocytes in the male marrow, along with an expansion of the more immature erythroid clusters. The female marrow showed the inverse trends for these clusters after LPS treatment. These data delineate distinct immune responses to LPS in erythroid cells based on maturation state and sex. In mammals, the effects of LPS on erythropoiesis are poorly defined. In a study conducted only in male mice, LPS suppresses the production of erythroid cells in the bone marrow via an indirect mechanism by acting on the erythroblastic island macrophage, a support cell for erythrocyte maturation^77,78^. Because erythropoiesis is suppressed in the bone marrow, the spleen readily takes over erythroid production in mice to replace cells lost to LPS-mediated hemolysis^79^. In zebrafish, erythroblastic island macrophages have yet to be characterized. Thus, changes in erythroid composition could also occur through alternative mechanisms, either indirectly through cytokine signaling from other immune cell types like those described in mammals or directly through an unknown pathway^80,81^. Additionally, histological analysis suggests that the zebrafish marrow is the primary site of erythroid generation, with the spleen functioning more as a reservoir and site of destruction for erythrocytes^82,83^. Similarly in humans, erythropoiesis does not usually switch from the bone marrow to the spleen and is therefore more functionally like zebrafish in some respects^84,85^. Thus, our data support the zebrafish marrow as a complementary animal model to study erythropoiesis and illuminates that the erythroid lineage in females responds uniquely to endotoxin-induced inflammation compared to males, an observation that has not been appreciated in any species due to the lack of female datasets.

Myeloid populations in the marrow were also highly dynamic in response to LPS-induced inflammation. Both sexes showed a reduction in the myeloid fraction following LPS exposure using flow cytometry analysis. These results recapitulate phenotypes in mice, where LPS treatment triggers rapid mobilization of neutrophils and monocytes from the bone marrow^52^. With scRNA-seq, we unmasked sex differences in the myeloid lineage; the abundance of nearly all macrophage and neutrophil clusters shifted in opposite directions between the sexes after LPS exposure. Moreover, we found that neutrophil composition changes varied with their maturation state. The less mature neutrophil subpopulations decreased in only the LPS-treated female, whereas the mature subpopulation decreased in only the LPS-treated male. A previous study has established that sex-biased responses to LPS in myeloid cell types can have an impact on survival^86^. In female mice, decreased numbers of neutrophils at the site of inflammation and increased serum levels of pro-inflammatory cytokines resulted in higher mortality than in males^86^. Mortality rates equalized between the sexes following the depletion of neutrophils, indicating that the sexual dimorphism in the neutrophil immune response was the driving factor in survival outcomes^86^. However, this study did not examine whether the disparate rates of neutrophil infiltration to the site of inflammation was due to sex differences in their mobilization from the marrow. Future studies are needed to determine how sex-dependent compositional changes in the marrow drive sexual dimorphic responses to inflammatory challenge in the periphery.

Male-female differences in the cellular makeup of the marrow are further underscored by the sex-biased transcriptional response to sustained LPS treatment in these lineages. Similar to the changes in composition, we observed DEGs in the scRNA-seq data for all hematopoietic cell types, with the greatest number of transcriptional differences occurring in the erythroid and myeloid clusters. Lymphoid clusters had comparatively modest changes to LPS exposure, which is likely attributed to the length of our experimental paradigm. Innate immune compartments are known to respond rapidly following LPS stimulation, whereas adaptive immune remodeling often occurs over longer timescales^87^. Therefore, chronic inflammatory models may reveal more pronounced sex-dependent effects within lymphoid populations.

Transcriptional sex differences persisted with sustained LPS exposure for many of the 37 genes that showed sex-specific response to acute LPS exposure. Several genes, including erythroid and myeloid differentiation factors and the well-known LPS-responsive cytokine *il1b* were only significantly changed in a particular cellular maturation state in one sex or the other. These results are consistent with a scRNA-seq study that observed sexual dimorphism in the transcriptomes of bone marrow-derived neutrophils of varying maturation states in mice^88^. Understanding the physiological significance of sex-biased immune responses for different maturation states is a clinically relevant avenue for future investigation as the underlying mechanisms could be leveraged therapeutically. While we are just beginning to catalogue all the compositional and transcriptional differences that exist between male and female immune cells of various maturation states, comparatively less is known about how these two factors interplay to produce a sex-dependent immune response. Our comprehensive dataset provides a valuable resource to the research community to untangle the unique compositional and transcriptional responses of hematopoiesis and immunity between the sexes. It is primed for mining the male- or female-specific transcriptional pathways that may underlie sex-specific protection or susceptibility.

An open question remains: What precisely is driving sex differences within the zebrafish marrow? Several of our datasets point to hormonal regulation. Firstly, we observed that marrow composition is dynamic in males and females throughout adulthood with the largest divergence between the sexes coinciding with peak reproductive maturity^48^—a process that requires regulation through sex hormone signaling^89^. In addition, a notable finding in this study was the conservation of sex-biased transcriptional programs in hormone signaling between zebrafish and humans. Despite experiencing ∼400 million years of evolutionary divergence^90^, both species shared several biological processes and regulators of hormone signaling via GO analysis and IPA of bulk RNA-seq datasets. Hormone-related functions were among the most significantly enriched GO terms for both sexes in zebrafish. In female marrows, these terms were specifically related to response to estrogen/estradiol. In male marrows, these terms were the broader categories of “response to hormone” and “steroid and cholesterol metabolic processes,” with cholesterol being an essential precursor for all steroid hormones. GO analysis of the human marrow DEGs revealed that metabolic and cholesterol-associated pathways were also among the top significant processes in human male marrow. Through IPA, we additionally showed that β-estradiol and progesterone are predicted upstream regulators in both zebrafish and human marrow datasets. Indeed, estrogen and progesterone are known to have immunomodulatory influences on HSPCs and differentiated blood cells in mammals^4,91,92^. Hormones have also begun emerging as a promising form of treatment for hematological diseases. Tamoxifen, a selective estrogen receptor modulator, has undergone clinical trials as a treatment for myeloproliferative neoplasms^93^, cancers that are known to have sex-biased subtypes and prognoses^94^. Taken together, the conservation of hormone-related pathways across species raises the possibility that sex-specific metabolic programs contribute to the regulation of hematopoiesis and immunity.

Our study establishes zebrafish as a model for investigating hematopoietic health and disease for both sexes, providing a complement to existing mammalian models. The mechanisms underlying sexual dimorphism have proven difficult to define in mammals due to contributions of both chromosomally-linked (XX/XY) gene dosage and gonadal hormone levels^19,95^. While genetic and surgical approaches have begun to disentangle chromosomal and endocrine influences^4,96^, their independent effects on hematopoiesis have yet to be fully resolved. A unique advantage of the zebrafish is that laboratory strains lack heteromorphic sex chromosomes^44,45^. This offers an exciting opportunity to specifically isolate the effects of hormones on the hematopoietic system, providing a better fundamental understanding of the mechanisms that underlie sex differences during aging and disease. Given the proven translatability of the zebrafish model^32,66^, it presents a highly scalable strategy to identify new therapeutic approaches for combating hematologic diseases.

## METHODS

### Zebrafish Husbandry

Adult zebrafish (*Danio rerio*) were maintained under standard laboratory conditions in a recirculating dechlorinated and filtered water system at 28°C under a 14-h light/10-h dark cycle. Embryonic and larval zebrafish were reared in 1x E3^97^ (pH 7.2) at 28°C then housed on the main system at 6 days post-fertilization (dpf). Zebrafish were fed a combination diet of GEMMA Micro (Skretting) and live artemia (Brine Shrimp Direct). The following zebrafish strains were used in this study: AB wild type strain and *Tg(cd41:GFP)*la2Tg^54^. All animal procedures were approved by the Institutional Animal Care and Use Committee (IACUC) at Dartmouth College (#00002271).

### Whole Kidney Marrow Dissociation

Zebrafish aged 2 months old through 24 months old were anesthetized in a lethal dose of buffered 0.04% Tricaine (pH 7) (MS-222; NDC: 50378-011-10; Syndel USA). Whole kidney marrow (WKM) dissection and single cell dissociation were performed as previously described with the following modifications^47^. Upon removal, WKMs were placed on ice in sterile Dulbecco’s phosphate buffered saline (dPBS; D8537; MilliporeSigma). WKMs were washed twice with dPBS and incubated with 30 uL of Liberase^TM^ (5401119001; MilliporeSigma) per 250 uL of dPBS and incubated at 28°C while shaking at 850 RPM for a total of 30 minutes. Once the tissue appeared fully disrupted, 600 uL of 1 mM EDTA/dPBS (pH 8.0) (BP24821; Fisher Scientific) was added and samples were incubated for 3 minutes at room temperature. Next, 200 uL of cold Fetal Bovine Serum (FBS) (A5670801; Gibco^TM^) was added, along with 1 mL of cold cell suspension media (0.8 mM CaCl2, 1x Penicillin-Streptomycin [15070063; Gibco^TM^], 0.5% FBS, Leibovitz’s L-15 Medium [21083027; Gibco^TM^]). Cells were pelleted at 300 RCF for 5 minutes at 4°C and washed twice with 1 mL of cold cell suspension media, pelleting each time. Cells were then resuspended in cold cell suspension media and strained through a 35 micron mesh filter into a sterile Eppendorf^TM^ tube. For cell viability analysis, cells were stained with DAPI (D9542; MilliporeSigma) at a concentration of 1:4000 then kept on ice.

### LPS Exposure

To induce acute inflammatory challenge, 6-month old zebrafish were anesthetized in 0.02% Tricaine (pH 7) (MS-222; NDC: 50378-011-10; Syndel USA) and administered intraperitoneal (IP) injections of either 5 mg/mL lipopolysaccharide (LPS; *E. coli* O111:B4; Cat.code: tlrl-3pelps; InvivoGen) or sterile phosphate buffered saline (PBS). Each fish received a total injection volume of 2 uL. For bulk RNA-sequencing or qRT-PCR, fish received a single IP injection of LPS or PBS and euthanized 2 hours post-injection. WKMs were rapidly dissected, flash-frozen in liquid nitrogen, and stored at - 80°C until RNA isolation. For flow cytometric analysis and single cell RNA-sequencing, fish received 2 IP injections of LPS or PBS, administered 24 hours apart. WKMs were immediately placed in dPBS and dissociated into single cell suspensions for downstream flow cytometric or single cell transcriptomic analysis.

### Flow Cytometry and Fluorescent Activated Cell Sorting (FACS)

Flow cytometry of single cell suspensions of zebrafish WKM was performed using a Beckman Coulter CytoFLEX flow cytometer. Fluorescence-minus-one controls were included where appropriate. Forward scatter area (FSC-A) and side scatter area (SSC-A) parameters were used to exclude debris. Live cells were identified by exclusion of DAPI. Doublets were removed by gating FSC-A by FSC-height (FSC-H). 30,000 live single cell events were collected per sample. Hematopoietic populations were defined based on previously established zebrafish WKM forward and side scatter characteristics corresponding to the erythroid, lymphoid, myeloid, and precursor fractions^27,47^. Data were analyzed using FlowJo^TM^ Software version 10.10.0 and gating strategies were applied consistently across all samples.

Fluorescent Activated Cell Sorting (FACS) was performed to exclude dead cells from the WKM single cell suspensions so that high purity live single cells could then be submitted for single cell RNA-sequencing. WKM single cell suspensions were sorted using a BD FACSAria cell sorter (BD Biosciences). Debris was excluded based on FSC-A and SSC-A, and doublets were removed by sequential gating of FSC-A by FSC-H and SSC-A by SSC-H. Sorting gates were applied consistently across all samples to ensure comparable cell populations were collected for downstream analyses. A total of 100,000 live single cells were sorted per sample and submitted for single cell RNA-sequencing library preparation.

### RNA Isolation

WKMs that were flash-frozen in liquid nitrogen were used for RNA isolation prior to qRT-PCR or bulk RNA-sequencing analysis. Total RNA was isolated using TRIzol Reagent (15596018; Invitrogen^TM^) according to manufacturer’s instructions. Briefly, frozen WKMs were homogenized in TRIzol, followed by phase separation with chloroform. The aqueous phase containing RNA was collected, precipitated in isopropanol overnight at - 20°C, and washed twice with 75% ethanol. RNA was resuspended in nuclease-free water and concentration and purity were determined using a NanoDrop spectrophotometer.

### qRT-PCR

For quantitative real-time PCR (qRT-PCR), cDNA was synthesized from 1 ug of RNA using the SuperScript™ IV First-Strand Synthesis System (18091300; Invitrogen^TM^) according to manufacturer’s instructions. qRT-PCR reactions were performed using KAPA SYBR FAST qPCR Master Mix (2x) (07959397001; Roche Diagnostics) on a CFX Opus 96 Real-Time PCR System (Bio-Rad). Genes interrogated in this study were *tnfa* (Fwd: 5’-GCTTCACGCTCCATAAGACCC-3’; Rev: 5’-AGAAGTGCTGTGGTCGTGTC-3’) and *il1b* (Fwd: 5’-ACGTCATCCAAGAGCGTGAA-3’; Rev: 5’-GTACGAGATGTGGAGCGGAG-3’), with *ef1a* (Fwd: 5’-TTCTGTTACCTGGCAAAGGG-3’; Rev: 5’-TTCAGTTTGTCCAACACCCA-3’) used as a housekeeping gene. Relative gene expression of *tnfa* and *il1b* were calculated using the comparative Ct (2^-ddCT) method, with expression normalized to *ef1a*.

### Bulk RNA Sequencing

RNA isolated from WKMs dissected from adult male and female 6 month old zebrafish exposed to either LPS (n = 5 males, n = 5 females) or PBS (n = 5 males, n = 5 females) were submitted for 3’ End-Counting sequencing (Illumina NextSeq 2000) at the Genomics Shared Resource at Dartmouth College. FASTQ reads were aligned to the Dr. Nathan Lawson Lab’s zebrafish transcriptome annotation reference^98^, quantified, and alignment metrics were assessed for quality control by the Genomic Data Science Core of Dartmouth College.

Healthy, adult human bone marrow bulk RNA sequencing data was selected from a previously published dataset^50^ (<u>Series GSE120444</u>) to compare to our zebrafish data. To minimize potential age-specific differences in the human dataset, male and female samples within +/-7 years of age were selected for analysis (Samples: J, U, B, H, O).

### Bulk RNA Sequencing Analysis

Raw, aligned counts from the adult zebrafish marrow were inputted into DESeq2 (version 1.46.0)^49^ to perform differential gene expression analysis with DESeq2’s standard workflow. Genes with at least 10 counts for the smallest group size (5 samples) were kept for analysis. An interaction term was added to the design formula (∼ sex + condition + sex:condition) in order to test for differences in response to LPS between the sexes. DESeq2 employs the Wald Test to calculate significance, and an adjusted p value (Benjamini-Hochberg procedure) < 0.1 was chosen as the cut-off for all bulk RNA sequencing differential gene expression comparisons.

A similar procedure was used for differential gene expression analysis of the human bone marrow selected from a previously published dataset^50^ (PMID: 30518681). In that case, genes with at least 10 counts for the smallest group size (2 samples) were kept for analysis, and the design formula tested only for differences between the sexes.

For gene ontology analysis of the biological processes in the zebrafish and human DEGs, sex-stratified lists of significant DEGs (468 zebrafish male DEGs, 605 zebrafish female DEGs, 166 human male DEGs, 209 human female DEGs) were uploaded to the “Functional Annotation” tool provided by DAVID Bioinformatics (NIH; https://davidbioinformatics.nih.gov). DAVID utilizes the Fisher’s Exact test to calculate significance, and a p value < 0.05 was chosen for the cut-off. We compared categories of significant DEGs in our zebrafish dataset to the human dataset.

### IPA Analysis

Sex-stratified zebrafish and human DEGs were also analyzed using Ingenuity Pathway Analysis^51^ (Qiagen Ingenuity® Systems, http://www.ingenuity.com). Input data included the PBS-treated male and female zebrafish marrow DEGs, converted to human orthologs for analysis (592 of 1,073 significant DEGs eligible for IPA), and DEGs from the curated male and female human marrow dataset (323 of 375 DEGs eligible for IPA) (**Supplementary Fig. 2a**). DEG eligibility is based on the genes present in the “Ingenuity Knowledge Base.” For “Biological Function” analysis, IPA maps inputted gene sets to a list of curated biological functions within their database and determines processes that are statistically enriched for both zebrafish and human DEGs. For “Upstream Regulator” analysis, IPA determines upstream molecules that are predicted to influence statistically enriched gene sets within the zebrafish and human DEG lists and determines shared pathways. IPA utilizes the Fisher’s Exact test to calculate significance for these analyses, and an adjusted p value (Benjamini-Hochberg procedure) < 0.1 was chosen for the cut-off.

### Single Cell RNA Sequencing

After FACS to obtain high purity live single cells, single cell suspensions of WKM from 6 month old zebrafish exposed to either LPS (n = 1 male, n = 1 female) or PBS (n = 1 male, n = 1 female) were submitted to the Genomics Shared Resource at Dartmouth College for Single Cell 3’ v4 (polyA) On-Chip Multiplexing (OCM) mRNA sequencing. Single cells were captured with the Chromium X instrument (10x Genomics). Libraries were prepped and sequencing was subsequently performed on the Illumina NextSeq 2000 instrument. Cell Ranger pipeline version 10.0.0 was used for processing and de-multiplexing raw sequencing data. FASTQ reads were mapped to the Lawson Lab’s zebrafish transcriptome annotation reference^98^ to generate single cell feature count matrices.

### Single Cell RNA Sequencing Analysis

Further processing of the gene count matrices generated in Cell Ranger was performed in R (version 4.4.1) with Seurat (version 5.5.0)^99^ and Tidyverse packages^100^. The matrices were filtered to remove genes that are expressed in fewer than 3 cells and remove cells that have fewer than 200 genes. Additional quality control steps removed cells with greater than 5% mitochondrial genes and fewer than 200 unique genes. To identify doublets, samples were preprocessed using the standard Seurat workflow as recommended by the DoubletFinder package^101^. DoubletFinder was employed to identify predicted doublets that were then removed from each sample. After this filtering, transcriptomes were acquired for 6990 PBS male cells, 9772 PBS female cells, 6889 LPS male cells, and 6001 LPS female cells. For further analysis, the four samples were merged into a single object of 29,652 cells.

The merged object was processed using the standard Seurat workflow, consisting of the following steps: Raw read counts were log-normalized using the “NormalizeData” function. 2000 highly variable features were identified using the “FindVariableFeatures” function, selected for using a variance stabilizing transformation (vst). The data was scaled with the “ScaleData” function to reduce skewing by highly-expressed genes. Linear dimensionality reduction was performed with a principal component analysis, using the function “RunPCA” and the list of variable features generated above. The top 35 principal components (PCs) which explained most of the variability in the data were identified based on PC heatmaps and PC ElbowPlot. These 35 PCs were used for clustering with the “FindNeighbors” function, and then cells were assigned to clusters with the “FindClusters” function. Finally, a UMAP was generated with the “FindUMAP” function using the same 35 PCs, and the resolution of the clusters was set to 0.775.

For downstream analysis, the “FindConservedMarkers” function was used to generate marker lists for each cluster. Analysis of the significant markers on these lists in combination with exploring expression distributions of dozens of known marker genes for hematopoietic, vascular, epithelial, and other kidney cell types, was used to annotate the clusters. The function “FindMarkers” was then utilized for differential gene expression analysis, comparing LPS vs PBS cells within the same cluster for males and females separately to determine LPS responsive genes. Significance of these DEGs was calculated using the Wilcoxon test and adjusted p value was found with the Benjamini-Hochberg procedure (cut-off set to p adjusted < 0.05). The gene patterns on the UMAP and violin plots were visualized with other Seurat functions.

## Data and Materials Availability

Bulk and single-cell RNA-seq data are submitted to the Gene Expression Omnibus GSE#### and GSE####.

## ACKNOWLEDGMENTS

We thank S. Niane and C. Mathews for fish husbandry and care. FACS was performed by the DartLab Immune Monitoring and Flow Cytometry Resource at Dartmouth College (RRID:SCR_019165) under the Dartmouth Cancer Center Support Grant P30CA023108. Bulk RNA sequencing was carried out in the Genomics and Molecular Biology Shared Resource (RRID:SCR_021293) at Dartmouth which is supported by NCI Cancer Center Support Grant 5P30CA023108 and NIH S10 (1S10OD030242) awards. Preprocessing and quality control steps were supported through Geisel School of Medicine at Dartmouth’s Center for Quantitative Biology through a grant from the National Institute of General Medical Sciences of the National Institutes of Health under Award Number P20GM130454. Single cell studies were conducted through the Dartmouth Center for Quantitative Biology in collaboration with the Genomics and Molecular Biology Shared Resource with support from NIGMS (P20GM130454) and NIH S10 (S10OD025235) awards.

## Author contributions

Experimental design and execution: S.E.P., C.L.W., E.A.J.; data analysis: S.E.P., C.L.W.; manuscript writing and editing: S.E.P., C.L.W., D.M.K.; supervision: D.M.K.

## Funding

This work was financially supported by grants from the National Heart, Lung, and Blood Institute (NHLBI; 1F32HL182156) awarded to S.E.P.; the National Institute of General Medical Sciences (NIGMS; 1R35GM160483) awarded to D.M.K.; and Developmental Funds from the “Friends of Dartmouth Cancer Center” (Dartmouth Geisel School of Medicine).

## Competing interests

The authors declare no competing interests.

## Notes

### Competing Interest Statement

The authors have declared no competing interest.

