## Supplementary for "Sexual dimorphism in adult zebrafish marrows regulates blood cell composition and immune response"

**This file includes:**

List of Supplementary Tables
Supplementary Figs. 1 – 5

**Other Supplementary Material for this manuscript include:**

Supplementary Tables 1 – 6

### List of Supplementary Tables

| | |
|---|---|
| **Supplementary Table 1** | Statistics for the changes of hematopoietic compositional flow cytometry gates throughout aging (**Supplementary Fig. 1**) |
| **Supplementary Table 2** | **Bulk RNA-seq DEGs** (**Fig. 2**)<br>Tab1: zebrafish PBS female vs PBS male marrow<br>Tab 1: human female vs male marrow |
| **Supplementary Table 3** | **Gene Ontology Terms for** (**Fig. 2**):<br>Tab 1: zebrafish male<br>Tab 2: zebrafish female<br>Tab 3: human male<br>Tab 4: human female |
| **Supplementary Table 4** | **LPS responsive DEGs** (**Fig. 4**):<br>Tab 1: In Venn diagram analysis<br>Tab 2: Sex-independent<br>Tab 3: Male<br>Tab 4: Female<br>Tab 5: Sex-dependent (*i.e.,* Interaction analysis) |
| **Supplementary Table 5** | Marker genes for single cell RNA-sequencing clusters used to annotate cell types (**Fig. 5**) |
| **Supplementary Table 6** | DEGs common between bulk RNA-seq interaction analysis and scRNA-seq clusters (**Fig. 6b**) |

### Supplementary Figures

#### Supplementary Fig. 1

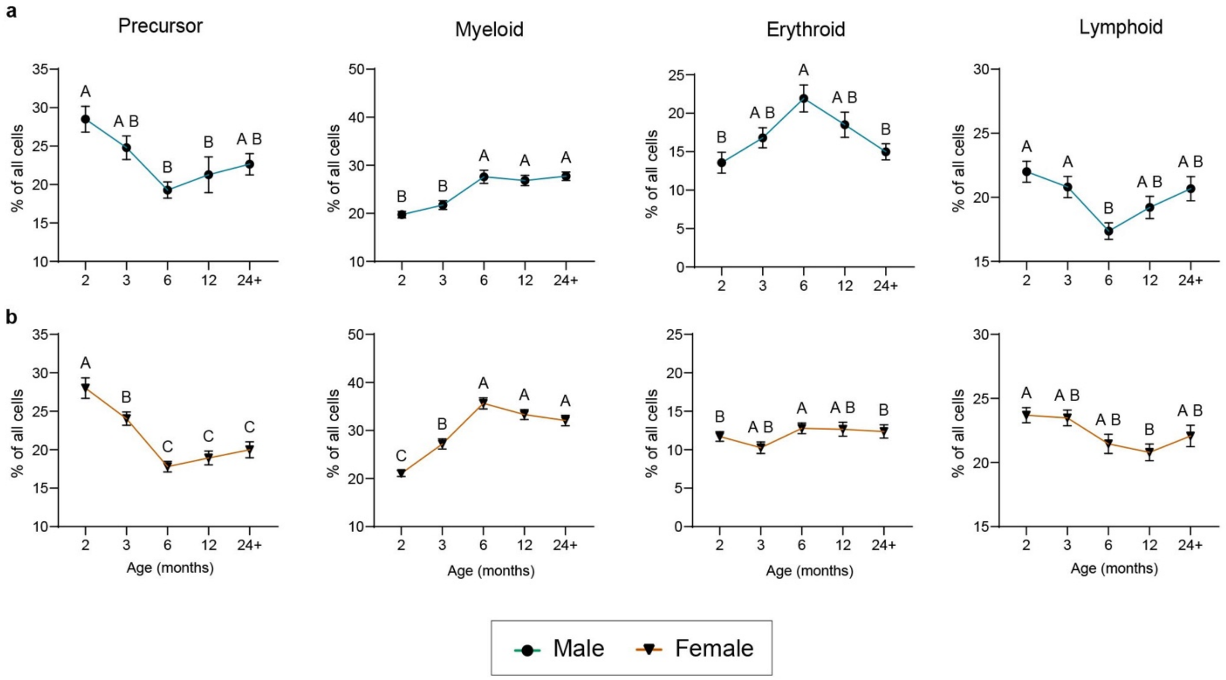


**Supplementary Fig. 1: Age is a determinant of blood cell diversity in zebrafish marrows.** (a-b) Percentage of live, single cells in each hematopoietic lineage gate for (a) male and (b) female marrows, aged 2 months (juvenile) through 24+ months (late adult). n = 15-20 marrows per group. pval <0.05 significance determined by an ordinary one-way ANOVA performed across ages for each sex, and is indicated through compact letter display. Data points with any shared letter are not significant, while data points with distinct letters are statistically significant. Error bars represent SEM.

**Supplementary Fig. 2**

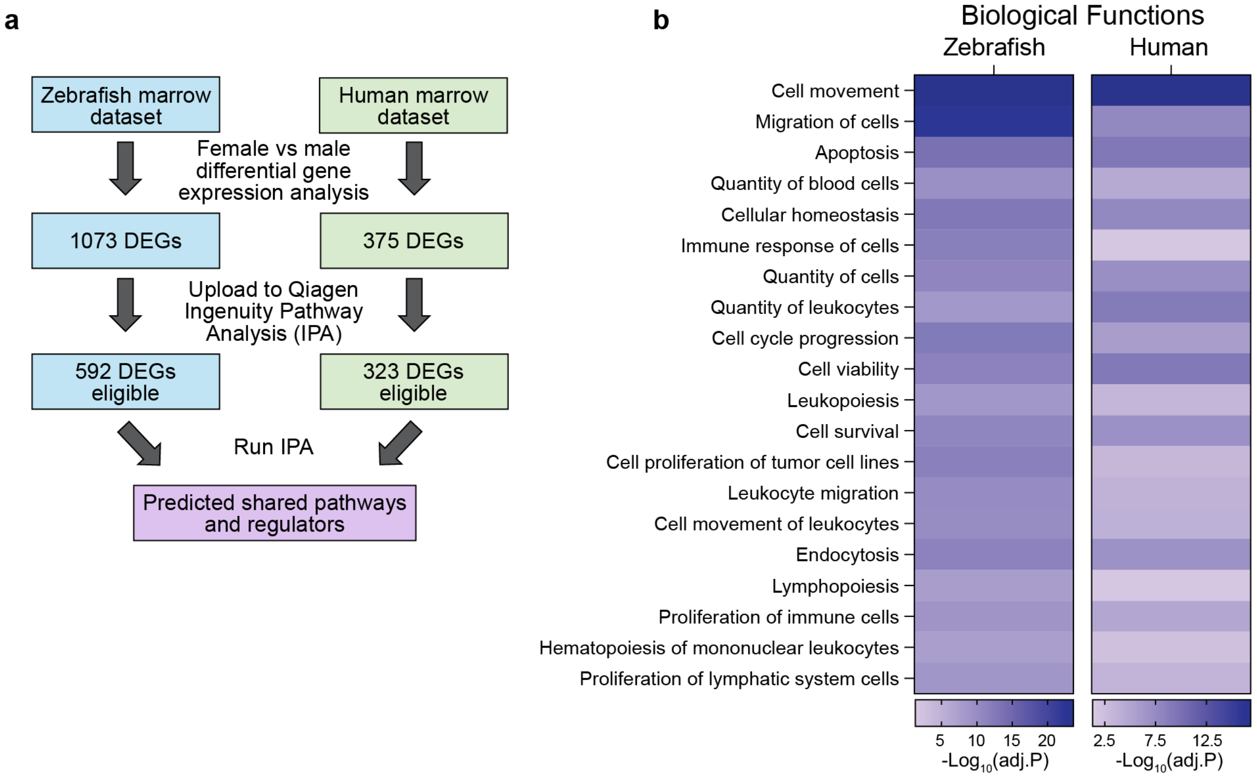


**Supplementary Fig. 2: Zebrafish and human marrow have conserved biological functions.** (a) Diagram of workflow for Ingenuity Pathway Analysis (IPA; Qiagen). DEG eligibility is based on the genes present in the "Ingenuity Knowledge Base." (b) Shared biological functions for zebrafish and human marrow from IPA, derived from bulk RNA-sequencing datasets (padj. <0.1, Fisher's Exact test).

**Supplementary Fig. 3**

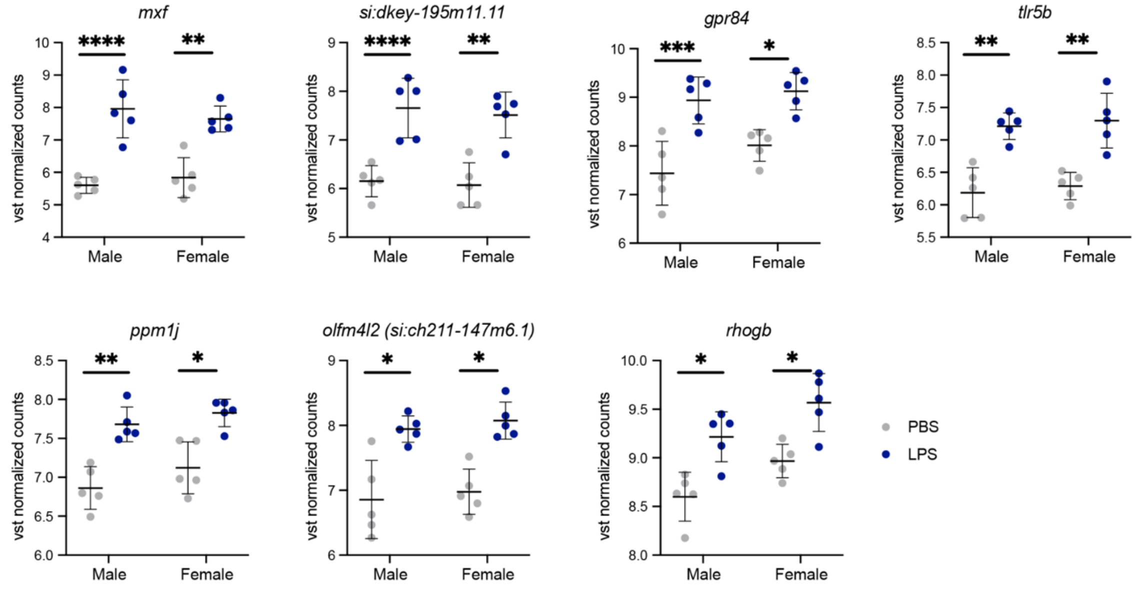


**Supplementary Fig. 3: Shared DEGs following LPS treatment.** Normalized transcript expression levels for the 7 shared LPS responsive genes in bulk RNA sequencing data (*mxf, si:dkey-195m11.11, tlr5b, si:ch211-147m6.1, gpr84, ppm1j, rhogb*) padj <0.1, Wald test.

### Supplementary Fig. 4

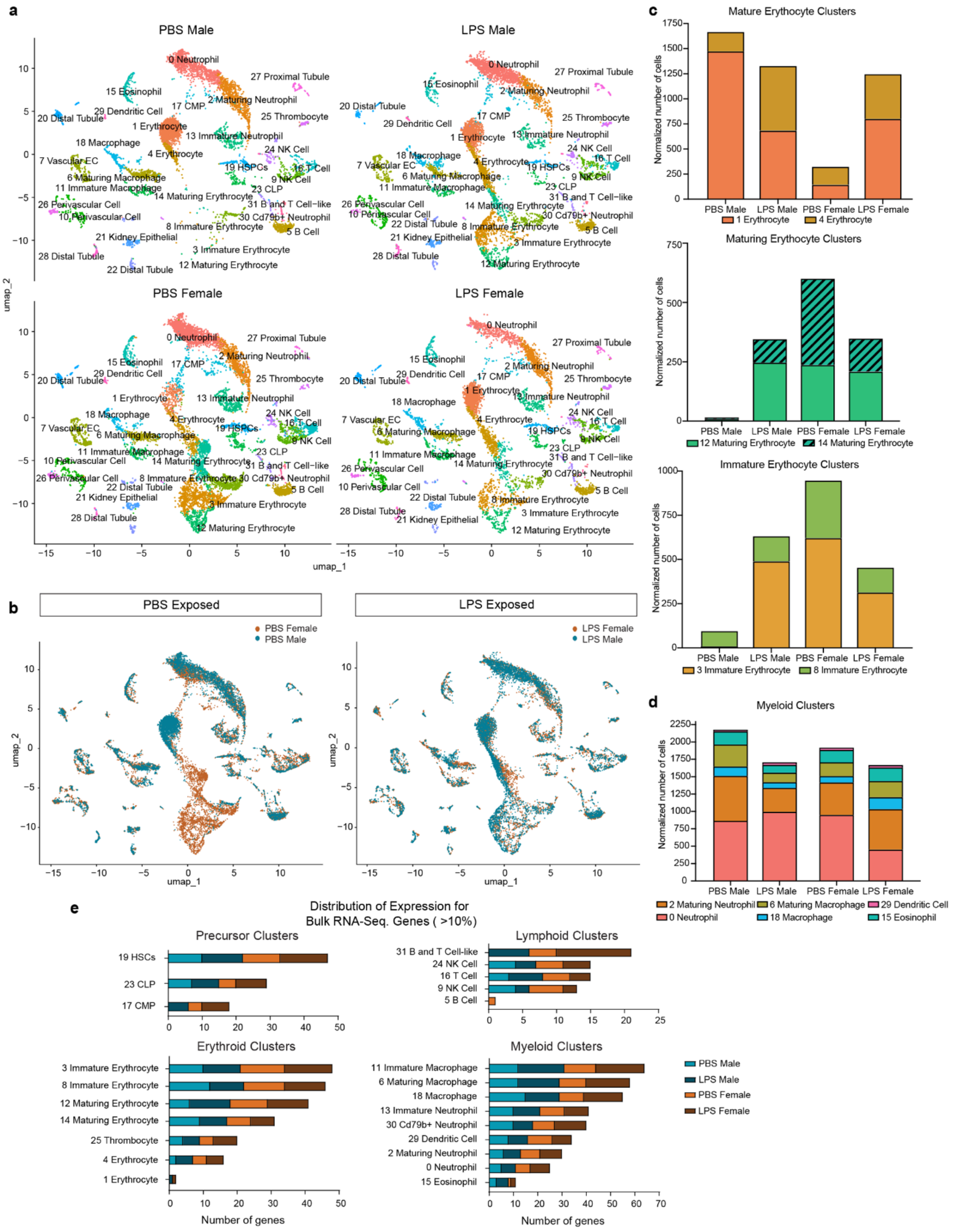

**Supplementary Fig. 4: Changes in cell proportions after LPS treatment.** (a) UMAP derived from scRNA-sequencing of 6-month-old male and female marrows from fish exposed to PBS or LPS. Fish were injected with PBS or LPS for 2 consecutive days and collected for scRNA-seq 24 hpi on the third day. (b) UMAP of male and female PBS or LPS exposed samples grouped by treatment. Orange and blue correspond to female and male samples, respectively. (c) Cell numbers for the mature erythrocyte, maturing erythrocyte, and immature erythrocyte clusters, normalized to the total number of all cells per sample. (d) Cell numbers for the more mature myeloid clusters, normalized to the total number of all cells per sample. (e) Distribution of expression of the significant DEGs from the bulk RNA-seq interaction analysis in the scRNA-seq clusters. 35 of the 37 bulk DEGs were identified in the scRNA-seq dataset. If a particular gene was expressed in >10% of cells in a given cluster, it was considered to be “expressed” in the cluster of that sample.

### Supplementary Fig. 5

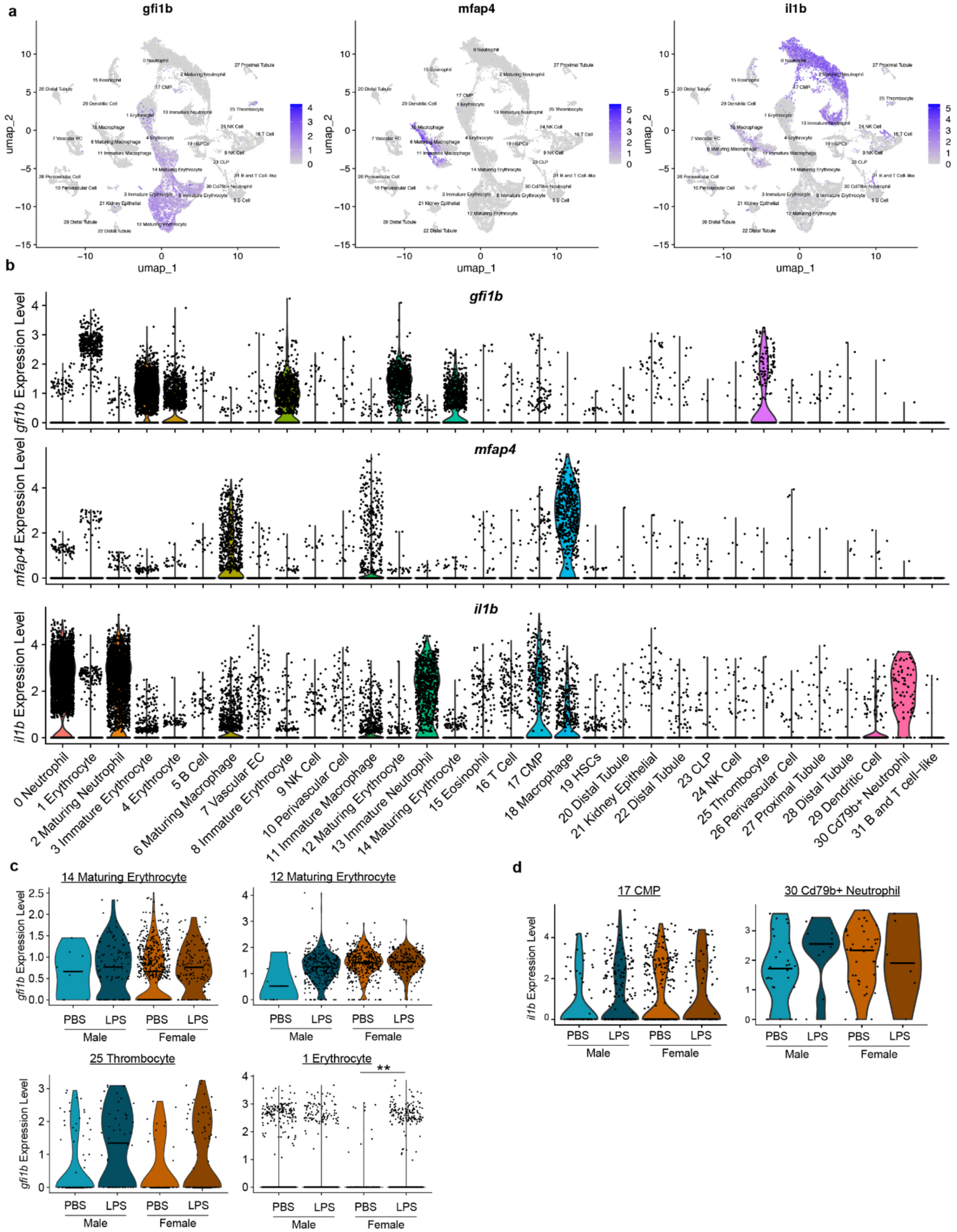

**Supplementary Fig. 5: Clusters expressing *mfap4*, *gfi1b*, and *il1b* transcripts.** (a) Feature maps highlighting clusters expressing *gfi1b*, *mfap4*, and *il1b*. (b) Violin plots of *gfi1b*, *mfap4*, and *il1b* expression in the scRNA-seq dataset. (c) *gfi1b* expression in PBS Male, LPS Male, PBS Female, and LPS Female samples in cluster 14 Maturing Erythrocyte, 12 Maturing Erythrocyte, 25 Thrombocyte, and 1 Erythrocyte. padj $<0.05$, Wilcoxon test used to calculate significance. (d) *il1b* expression in PBS Male, LPS Male, PBS Female, and LPS Female samples in cluster 17 CMP and 30 Cd79b+ Neutrophil.
